# Autonomic arousal tracks outcome salience not valence in monkeys making social decisions

**DOI:** 10.1101/2020.02.17.952986

**Authors:** Benjamin M. Basile, Jessica A. Joiner, Olga Dal Monte, Nicholas A. Fagan, Chloe L. Karaskiewicz, Daniel R. Lucas, Steve W. C. Chang, Elisabeth A. Murray

**Affiliations:** Section on the Neurobiology of Learning and Memory, Laboratory of Neuropsychology, National Institute of Mental Health, National Institutes of Health, Bethesda, MD, 20892, USA; Department of Psychology, Yale University, New Haven, CT, 06511, USA; Department of Psychology, University of Turin, Torino, Italy; Department of Neuroscience, Yale School of Medicine, New Haven, CT, 06520, USA; Kavli Institute for Neuroscience, Yale School of Medicine, New Haven, CT, 06520, USA

**Keywords:** vicarious reinforcement, social valuation, pupillometry, rhesus monkeys, anterior cingulate cortex, prosociality

## Abstract

The evolutionary and neural underpinnings of human prosociality are still largely unknown. A growing body of evidence suggests that some species find the sight of another individual receiving a reward reinforcing, often called vicarious reinforcement. One hypothesis is that vicarious reward is reinforcing because it is arousing like a primary reward. We evaluated this hypothesis by measuring the autonomic pupil response of eight monkeys across two laboratories in two different versions of a vicarious reinforcement paradigm. Monkeys were cued as to whether an upcoming reward would be delivered to them, another monkey, or nobody and could accept or decline the offer. As expected, all monkeys in both laboratories showed a marked preference for juice to the self, together with a reliable prosocial preference for juice to a social partner compared to juice to nobody. However, contrary to the autonomic arousal hypothesis, we found that pupils were widest in anticipation of juice to the self, moderately-sized in anticipation of juice to nobody, and narrowest in anticipation of juice to a social partner. This effect was seen across both laboratories and regardless of specific task parameters. The seemingly paradoxical pupil effect can be explained by a model in which pupil size tracks outcome salience, prosocial tendencies track outcome valence, and the relation between salience and valence is U-shaped.

## 1. Introduction

Humans watch game shows partly because we like seeing others get rewarded. This phenomenon is often called vicarious reinforcement. A growing body of comparative evidence suggests that vicarious reinforcement is a fundamental cognitive mechanism supporting social behavior in primates. For example, rhesus monkeys will choose to give juice to a partner monkey more often than choose to withhold juice^1^. Monkeys that choose to give juice to another monkey also work to withhold aversive air puffs from that same monkey and these choices correlate with the strength of the pair’s affiliative relationship^2^. Moreover, chimpanzees will choose to deliver rewards to both themselves and another chimp over just themselves^3^. Importantly, the tendency to give reward in these experimental settings depends on the presence of the other individual; no prosocial tendency is shown when reward goes to a collection jar instead of a conspecific. However, prosocial behaviors are not always the prepotent tendency in primates. Monkeys tend to defect rather than cooperate in classic economics games^4^, offering reward to another monkey can cause monkeys to work less^5^, the same monkeys who choose to give juice to another rather than have it go to nobody will also choose to only get juice themselves rather than get juice jointly with another monkey^1^, and both monkeys and apes often show robust disregard across multiple tasks for whether a partner receives a reward^6^. Thus, it is still unclear what features and parameters modulate vicarious reinforcement and how much it generalizes to different situations.

Researchers have made good progress in understanding the cognitive and neural underpinnings of vicarious reinforcement via studies of monkeys performing social reward allocation tasks. In one prominent example of this task^1^, two monkeys – an actor and a recipient – sit at right angles to each other. Each faces a computer screen that displays visual cues that predict juice (reward) outcomes. The actor monkey is either cued about an upcoming juice outcome or chooses between two outcomes. The typical outcome conditions are juice to the *self*, juice to the *other* monkey, juice to *both* monkeys, or juice to *neither* monkey. Critically, these options are always paired in choice trials such that there is no primary reward gain or loss from the perspective of the actor monkeys, controlling for a confound in self reward contingency. As expected, monkeys strongly prefer receiving juice themselves. Interestingly, they also prefer juice being received by the *other* monkey over *neither* monkey. In this behavioral paradigm, neurons in the rostral anterior cingulate gyrus (ACCg) code the chosen social outcome^7^ and neurons in the amygdala code the value of juice amount similarly regardless of whether it is delivered to the self or the other monkey, but not when it is delivered to a jar in the nonsocial control condition^8^. In a similar paradigm, researchers found neurons in the dorsal convexity of the medial prefrontal cortex that selectively coded reward for either the actor monkey or a partner monkey^9^.

One hypothesis for the vicarious reinforcement effect is that monkeys’ prosocial tendencies are based on the autonomic arousal associated with anticipation of the reward outcome. Accordingly, they choose reward to the *self* most often because it is most arousing, reward to *other* moderately often because it is moderately arousing, and reward to *neither* least often because it is least arousing. This would be consistent with how monkeys’ pupil size, a common indicator of autonomic arousal, behaves during nonsocial tasks: pupil dilation reliably increases with the amount of juice predicted by a stimulus^10^. Neurally, it would be consistent with the population average activity of ACCg neurons; these neurons are most active for rewards to the *self*, moderately active for rewards to the *other*, and least active for rewards to *neither*^7^.

To evaluate the degree to which monkeys’ prosocial tendencies are linked to their arousal, and thus guide future research in investigating neural computations guiding these social judgments, we measured pupil size as monkeys chose whether to accept or reject juice offers to themselves or a partner in a social reward allocation task. If social preferences are driven by arousal, then we predict that pupil size will scale monotonically with prosocial tendencies, with pupil largest in anticipation of *self* rewards, next largest in anticipation of *other* rewards, and smallest in anticipation of *neither* rewards. If this pattern of pupil size is not found, then some other factor must be responsible for prosocial tendencies. To assess the generality of our findings, we conducted this study in two separate laboratories that used monkeys with different life histories, behavioral test setups with different physical arrangements, stimuli with different perceptual properties, and social reward allocation tasks with different parameters. Experiment 1 reports results from the laboratory in Bethesda, MD and Experiment 2 reports results from the laboratory in New Haven, CT.

## 2. Experiment 1 – Bethesda laboratory

### 2.1 Methods

#### 2.1.1 Subjects

Nine adult male rhesus macaques (*Macaca mulatta*) housed at the National Institute of Mental Health in Bethesda, MD participated in the experiment (mean age at start = 6.5 yrs), six as actor monkeys and three as recipient monkeys. Monkeys were housed singly due to the constraints of a subsequent experiment, but had visual and auditory access to multiple conspecifics in the room. Two actors each were assigned to a dedicated recipient and housed directly across from that recipient. Thus, all actors and recipients were familiar with each other. Housing was on a 12:12 light:dark cycle with *ad libitum* food. Daily fluid was controlled such that monkeys maintained good test motivation in the test apparatus, good health, and a weight above 85% of their free-feeding weight. Prior to this study, we implanted each monkey with a titanium head post to allow head-restrained eye tracking ^11^ and shaped each monkey to perform a basic oculomotor saccade task. All procedures were reviewed and approved by the National Institute of Mental Health (NIMH) Animal Care and Use Committee and complied with US law.

#### 2.1.2 Apparatus and Stimuli

We tested monkeys in pairs in a sound-attenuating chamber (Crist). Actors sat in a primate chair facing a computer monitor (22.86 cm wide × 30.48 cm tall) at a distance of approximately 54 cm. Recipients sat in a primate chair such that their head was immediately to the right of the monitor (actor’s view) and they faced over the actor’s right shoulder (Fig 1a). This placed both monkeys in easy view of each other, but not directly facing each other as direct gaze can evoke aggression in rhesus macaques^12^. Both monkeys were head restrained during testing. A camera positioned at the lower right corner of the monitor tracked the actor’s eye position and pupil width. Juice (50:50 apple juice:water) was delivered via hidden tubing to one of two metal spouts positioned at the mouth of either the actor or recipient. Pressurized juice-delivery systems (Precision Engineering) were housed outside the chamber and delivery was gated by solenoids housed in their own sound-attenuating box. This box effectively silenced the juice delivery, rendering it undetectable by two separate humans who performed forced-choice and yes-no detection tests (proportion correct = 50% and d’ = 0.0, respectively). In addition, a sound meter placed ∼5 cm away from the box did not register any sound increase from rapid solenoid firing when the lid was closed (max. sound level during juice delivery with sound-attenuating box open = 58.82 db (± 0.60), during delivery with box closed = 49.89 db (± 1.10), and not during delivery = 50.42 db (± 1.17)). Still, to rule out any contribution of the solenoid to monkey’s behavior, we took two additional precautions. First, the sound-attenuating box housed a third dummy solenoid that fired on *neither* reward trials, and a recorded audio clip of a solenoid firing was played inside the monkey testing chamber on every completed trial regardless of reward outcome. Stimuli were two abstract shapes that could appear in one of four orientations to signal the start of the trial or one of the three juice offers (Fig 1c). One shape was used on Social sessions and the other was used on Nonsocial control sessions in which the recipient monkey was replaced with a juice collection receptacle (Fig 1b).

**Figure 1.**
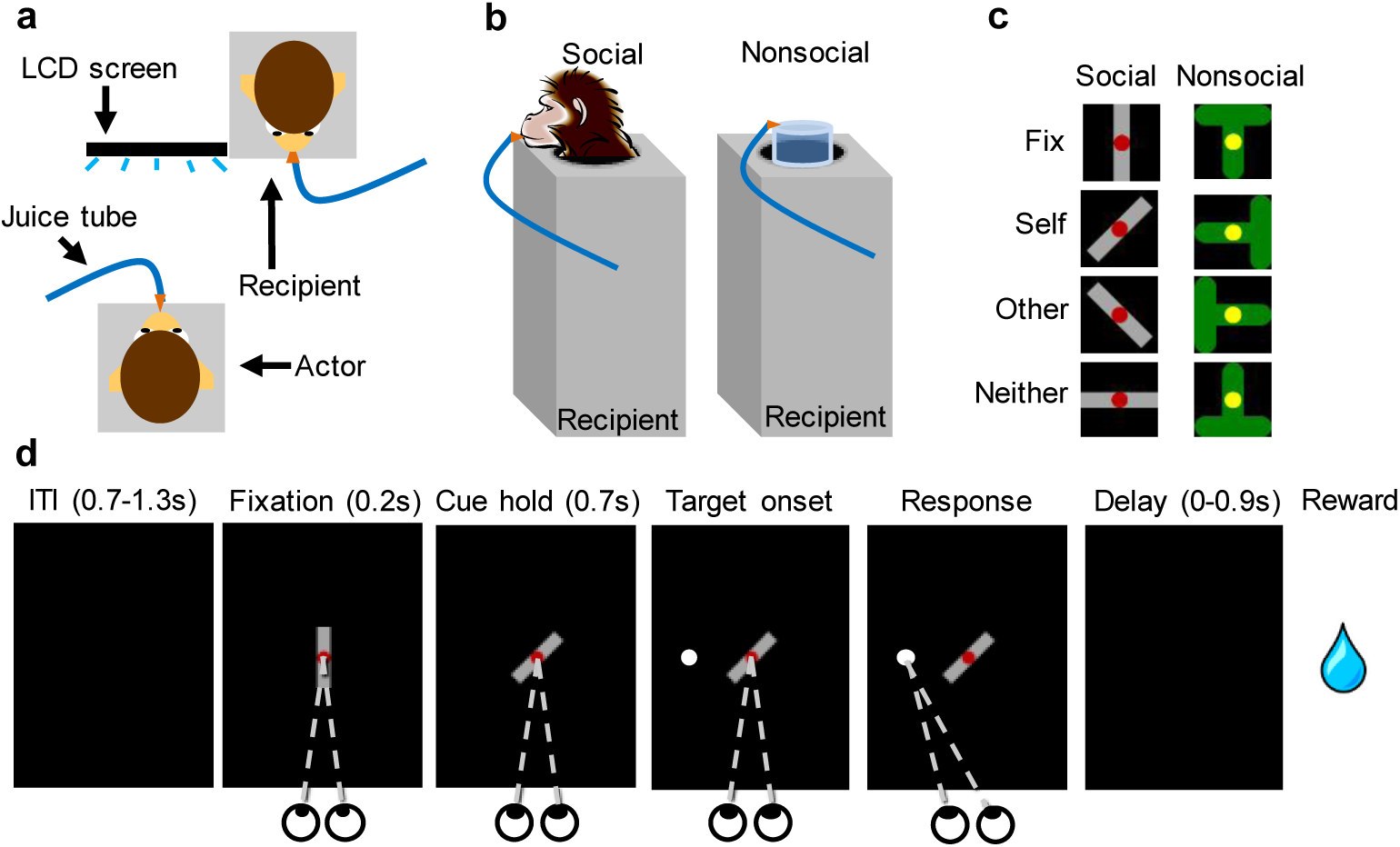
Social reward allocation task used in Experiment 1. a. Top-down schematic of the test arrangement with the actor monkey facing an LCD screen next to a recipient. b. Schematic side view of juice delivery to recipient or juice collection cylinder in Social and Nonsocial sessions. c. Stimuli used in the Social and Nonsocial sessions. The cues used for fixation were rotated to create the three reward conditions. d. Schematic of the trial progression in a Social session in which the stimulus signals that ‘reward to self’ is on offer. Each square depicts the LCD screen as seen by the monkey. If the monkey completed the saccade to the peripheral target, the reward condition on offer for that trial was implemented. Note that the white peripheral saccade target appeared equally often in one of eight locations equidistant from the center.

#### 2.1.3 Behavioral procedures

Two monkeys participated in the task at a given time, one actor and one recipient. The six actors were matched with three dedicated recipients such that each recipient worked with two actors, actors always worked with the same recipient, and no actor ever served as recipient.

Each trial began with the onset of the fixation stimulus (Fig 1d). After an actor monkey acquired and held central fixation for 0.2 s, the stimulus was replaced with one of three alternative orientations that predicted one of three juice outcomes: *self, other*, or *neither. Self* trials delivered juice to the actor, *other* trials delivered juice to the recipient on Social sessions or the juice receptacle on Nonsocial sessions, and *neither* trials delivered no juice. To accept the juice offer, the actor monkey had to maintain fixation for an additional 0.7 s until a peripheral saccade target appeared in one of eight equidistant locations, and then had to make a saccade to that target. After a random delay of 0.0-0.9 s, the signaled juice outcome was delivered, and the actor had an additional 1 s of free viewing time to observe the recipient. To reject the juice offer, the actor could abort fixation after the rotated cue appeared or fail to saccade to the peripheral target. Aborted trials were followed by a white screen that lasted 5 s and were repeated if the actor aborted before having seen the juice offer but not repeated if the actor had seen the juice offer. All trials were separated by a blank interval of 0.7-1.3 s. Actors worked for either 0.3 or 0.5 ml of juice per reward, depending on individual motivation, and recipients always received 0.5 ml of juice per reward. The amount of juice per reward was held constant within a given session. The delivery times were calibrated such that juice delivery, or unfilled interval if it was a *neither* offer, lasted the same duration for all three conditions. Juice offers were pseudo-randomly determined, with the constraints that half of offers were *self* to maintain motivation, there were an equal number of *other* and *neither* offers, and an offer could appear no more than four times in a row. Monkeys completed one 600-trial session per day. Nonsocial sessions were identical to Social sessions except for the use of a different stimulus and the presence of a juice receptacle instead of the recipient monkey. Social and Nonsocial sessions were run in blocks of 10 sessions with an ABBA or ABAB pattern, with half of monkeys assigned to each pattern.

#### 2.1.4 Data analysis

Completion rates of *other* and *neither* trials were compared using paired t tests. We analyzed both as a group across individuals and for each individual monkey across sessions. Pupil traces were smoothed with a zero-phase low-pass digital filter using the filtfilt function in MatLab (MathWorks, Inc.) to compensate for the fact that our data acquisition system records at higher frequency than is sent by the eye tracker. Outliers in which the value at a particular millisecond was more than 3 SD away from the median of all other trials of that same type in that session were removed. We normalized the data for each trial as a proportion change from the initial 50 ms of that trial during fixation. All pupil data were expressed as z values, as in previous investigations of pupil size^13,14^ to control for individual differences in pupil dynamic range. Statistical analyses were run on the last 50 ms of fixation to the cue and on the 50 ms of hold on the peripheral saccade target. Trial completion rates and pupillary changes were analyzed via two-way ANOVA with outcome and session type (Social and Nonsocial) as factors. All tests were two tailed with an alpha of 0.05. Four actors completed 20 sessions each of Social and Nonsocial trials, one actor completed 40 sessions of each type, and one actor completed 50 sessions of each time. To ensure the same amount of data was analyzed for each animal regardless of learning rate, data analysis was limited to the last 20 Social sessions and the last 20 Nonsocial sessions. This number of sessions is similar to that reported in Experiment 2.

### 2.2 Results

In the Social sessions, monkeys completed the most *self* trials, the next most *other* trials, and the fewest *neither* trials (Fig 2a). There was an interaction between outcome and session type (F_(2,10)_ = 4.84, p = .034; partial η^2^ = .492) illustrating that trial completion rates depended on both the juice offer and whether the partner was present. The critical preference for *other* trials over *neither* trials was significant at both the group level (t_5_ = 5.87, p = .002, d = 2.40) and for each of the six individual monkeys (all ps < .028). In the Nonsocial sessions, during which the recipient partner (*other*) was replaced with a juice collection cylinder, there was no preference for *other* over *neither* trials (Fig 2c; t_5_ = 1.27, p = .260). This reproduces the main behavioral finding from Chang et al. ^1,7,8,15^, showing a reliable prosocial preference for giving juice to another monkey over wasting juice. Further, it demonstrates that the effect depended on the presence of the other monkey.

**Figure 2.**
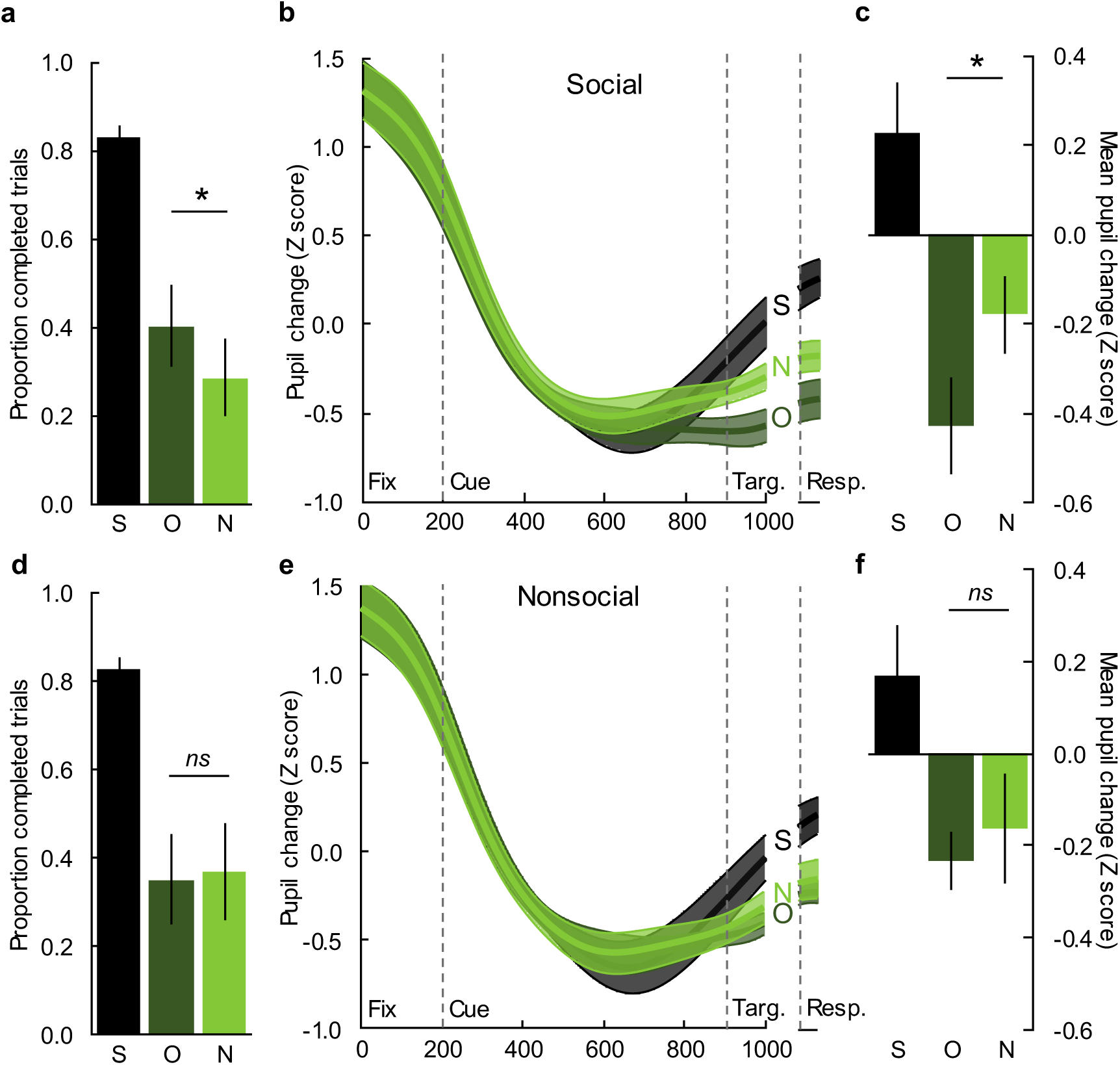
Pupils were more constricted in anticipation of preferred prosocial *other* trials than *neither* trials. a. Proportion (±SEM) of *self* (S), *other* (O), and *neither* (N) trials completed in the Social sessions. b. Relative pupil change across the trial in the Social sessions aligned to the onset of fixation (Fix) and to the saccade response (Resp). Error bars and shaded bands are ±SEM. c. Mean (±SEM) pupil diameter for each reward outcome in Social sessions during fixation on the peripheral target, before reward delivery. d e & f. Same as for a-c, above, but for the Nonsocial sessions.

Monkeys’ pupils constricted in the first half of the trial with the increased light from the fixation stimulus and then rebounded in the second half of the trial in anticipation of the reward outcome (Fig 1b & d). In Social sessions, this rebound was largest in anticipation of reward to *self*, moderate in anticipation of reward to *neither*, and, surprisingly, least in response to reward to *other* (Fig 2b). This difference was significant both in the epoch just before breaking central fixation and the epoch fixating on the peripheral saccade target before reward delivery (central fix: t_5_ = 3.13, p = .026, d = 1.28; peripheral fix: t_5_ = 3.47, p = .018, d = 1.42). Notably, the ordering of the pupil effect, *self*>*neither*>*other* was different than the ordering of the trial completion effect, *self*>*other*>*neither*. In the Nonsocial control sessions, pupil size was still widest in anticipation of *self* rewards, but did not differ between *other* and *neither* trials (Fig 2d; central fix: t_5_ = 0.98, p = .372; peripheral fix: t_5_ = 0.90, p = .410). A two-way ANOVA with session type and outcome as factors found an interaction (F_(2,10)_ = 4.32, p = .044; partial η^2^ = .464), where pupil diameter differences between outcome conditions depended on session type. This demonstrates that the pupil size difference between *other* and *neither* trials, like the trial completion rates, depended on the presence of the recipient monkey.

## 3. Experiment 2 – New Haven laboratory

### 3.1 Methods

#### 3.1.1 Subjects

Four rhesus macaques housed at Yale University in New Haven, CT, two males (monkeys K and H) and two females (monkeys E and C), aged 6-12 years, participated in this experiment. Monkeys were socially housed in pairs but were not matched with cage mates during the experiments. However, all four participating monkeys had visual access to one another in the colony room. Housing was on a 12:12 light:dark cycle with *ad libitum* food. Daily fluid was controlled such that monkeys maintained both good motivation in the test apparatus, good health, and a weight above 85% of their free-feeding weight. Prior to this study, we implanted each monkey with a head post (Crist Instruments or GrayMatter Research) to allow head-restrained eye tracking and shaped each monkey to perform a basic oculomotor saccade task. All procedures were conducted in accordance with the *Guide for the Care and Use of Laboratory Animals*^16^ and with approval from the Yale University Institutional Animal Care and Use Committee.

#### 3.1.2 Apparatus and stimuli

Each monkey faced its own display screen; the screens were situated at a 90° angle from one another. The recipient monkey was always situated diagonally across from the actor monkey to the right from the actor’s screen (Fig 3). Each monkey was fitted with a juice tube for delivering rewards. The solenoid valves that delivered the liquid rewards were placed in another room to prevent monkeys from forming secondary associations between solenoid clicks and different reward types. Three separate solenoids were used for delivering juice to the actor (*self*), the recipient (*other*), and to the juice collection bottle (*neither)*, thus controlling for secondary associations. All experiments were carried out in a dimly lit room to ensure visibility of the actor and recipient monkey. Both actor and recipient were head-restrained during the experiments. Eye position and pupil diameter were recorded at 1,000 Hz (EyeLink, SR Research). Stimuli were colored squares. Different colors signaled different reward conditions. Stimuli were controlled by a computer running custom software (Picto).

**Figure 3.**
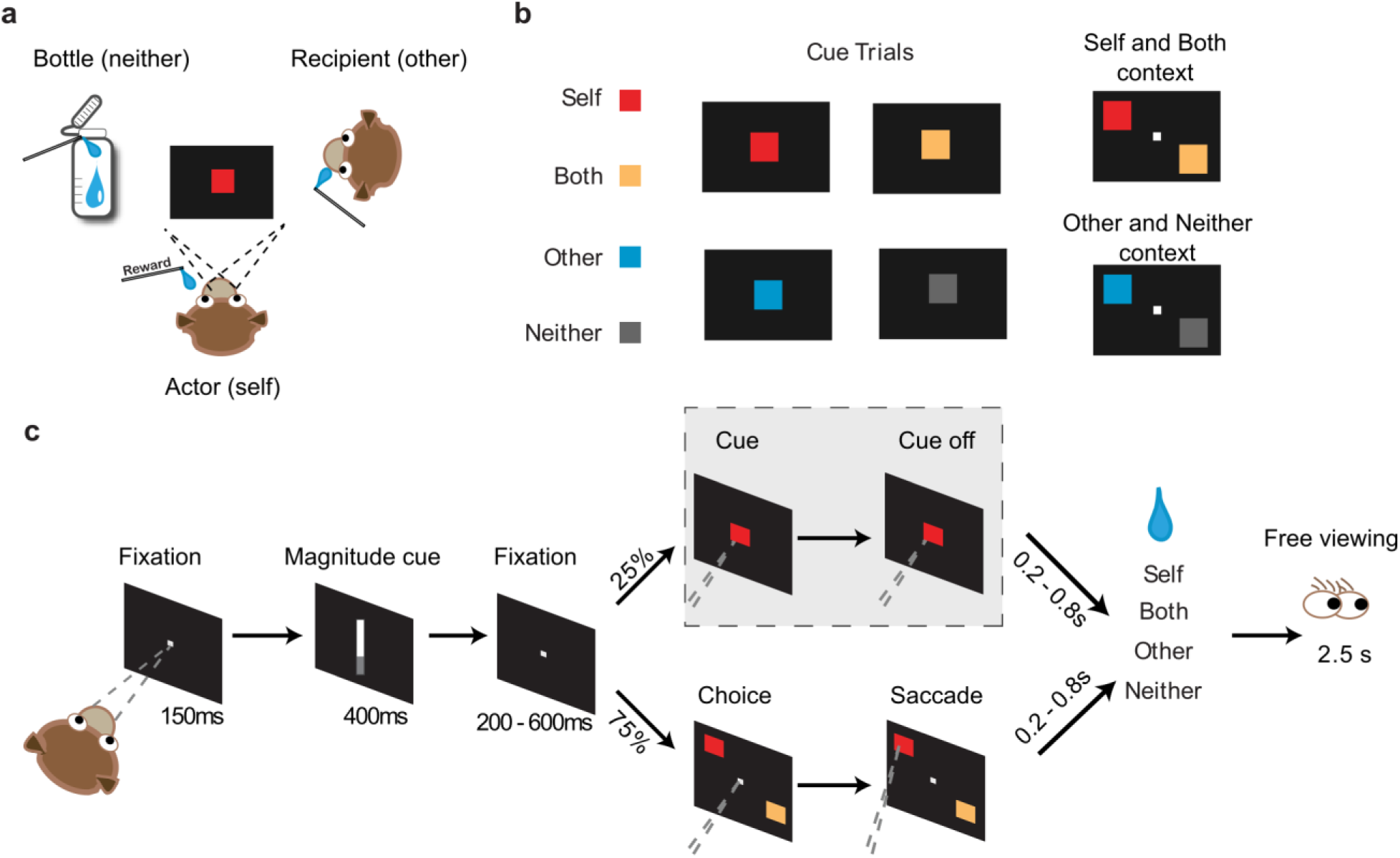
Social reward allocation task used in Experiment 2. a. Top down schematic of the testing arrangement with the actor monkey facing an LCD screen next to a recipient and an empty juice collection bottle. The recipient also faced his own LCD screen (not pictured), which showed the same stimuli. b. Left, example stimuli on the cued trials, in which a reward predicting cue that appeared on the center of the screen mapped onto a juice reward delivered to the actor (*self*), the recipient (*other*), both monkeys (*both*), or the juice collection bottle (*neither*). Right, example stimuli on choice trials, in which the actor chose between delivering a reward to self or to self and the recipient (*self* vs. *both*) on some trials and between delivering a reward to the recipient or the bottle (*other* vs. *neither*) on other trials. c. A schematic of the trial progression.

#### 3.1.3 Behavioral Procedures

Two monkeys participated in the task at a given time, one actor and one recipient. Monkeys K and H (males) played the role of actor, while monkeys E and C (females) played recipient to K and H, respectively.

An actor began a trial by fixating on a central square for 150 ms. The reward value on each trial was then specified by a vertical bar indicating juice volume (0.2, 0.4, or 0.6 ml). The actor was required to maintain fixation on the vertical bar for 400 ms. Following a variable delay (200, 400, or 600 ms), the actor was presented with either a choice (75%) or a cued (25%) trial. On cued trials, a cue signaling reward outcome (*self, other, both*, or *neither*) was presented at the center of the screen. To accept the offer, the actor had to maintain fixation for 150 ms. Upon successful completion of the fixation requirement, there was a random delay (200, 400, 600, or 800 ms) before the cued juice outcome was delivered to the actor (*self* cue), the recipient (*other* cue), both the actor and the recipient (*both* cue), or no one (*neither* cue). After the reward delivery, the actor had an additional 2.5 s of free viewing time during which he was free to look at the recipient or any other locations in the setup. To reject the juice offer, the actor could simply abort fixation after the shape cue appeared. Aborted trials were followed by a white screen that lasted 5 s and were repeated if the actor aborted before having seen the juice offer but not repeated if the actor had seen the juice offer. All trials were separated by a blank interval of 2.5 s. On choice trials, two cues appeared on the screen simultaneously, one to each side of the center. To ensure the actors had nothing to gain or lose with respect their own reward outcome, there were only two possible choices on offer: *self* vs. *both* and *other* vs. *neither*. Again, these options were always paired in choice trials such that there is no primary reward gain or loss from the perspective of the actor monkeys, controlling for a confound in self reward contingency. Timing of choice trials was identical to that of cued trials except now monkeys needed to make a saccade to select their choice.

Cued and choice trials were pseudo-randomly interleaved. As in Experiment 1, juice offers (*self, both, other*, and *neither*) were pseudo-randomly determined on the cued trials, with equal probabilities. On *both* trials (cued and choice), the two monkeys received the same amount of juice at the same time. The *neither* trial delivered juice to a bottle situated across from the recipient monkey, to the left of the actor. Combining both monkeys, 57 days of data were collected with 315.75 ± 119.11 (M ± SD) trials per day.

#### 3.1.4 Data Analysis

Data from the choice trials were used to evaluate each monkey’s social preferences. Only completed choice trials were included. Preference was measured via proportion of each choice for the two trial types. Differences in proportion choice were analyzed using a t test.

Data from the cued trials were used to determine pupil responses to avoid the potential confound associated with measuring pupil diameter on choice trials involving eye movements. Only completed cue trials were analyzed. Data were smoothed with a zero-phase low-pass digital filter using the filtfilt function in Matlab. Pupil diameter was normalized trial by trial to the 150 ms fixation period. All pupil data were expressed as z values, as in previous investigations of pupil size^13,14^, to control for individual differences in pupil dynamic range. Pupil data were analyzed across the four outcomes from 200-800 ms after cue onset using a one-way ANOVA and post hoc Tukey test. Analysis of additional epochs yielded similar results. All tests were two tailed with an alpha of 0.05.

### 3.2 Results

Previously, we have shown that actor monkeys develop a typical pattern of social preferences in the social reward allocation task; they choose *other* over *neither* (prosocial preference), and choose *self* over *both* (antisocial preference)^1,7,8,15^. Here, we first replicated this behavioral finding (Fig 4a). Both actor monkeys significantly preferred choosing *self* (M ± SEM; 0.55 ± 0.01,) over *both* (0.45 ± 0.01) reward outcome (t_96_ = 6.01, p < 0.001). This is consistent with previous work showing monkeys to be antisocial in reward contexts where they themselves receive a reward. Critically, monkeys preferred choosing *other* (0.67 ± 0.01) over *neither* (0.33 ± 0.01) reward outcome (t_96_ = 25.20, p < 0.001) indicating a prosocial preference in the *other* versus *neither* trials when they themselves could not receive a reward. This is consistent with monkeys having context-dependent prosocial and antisocial preferences in the social reward allocation task.

**Figure 4.**
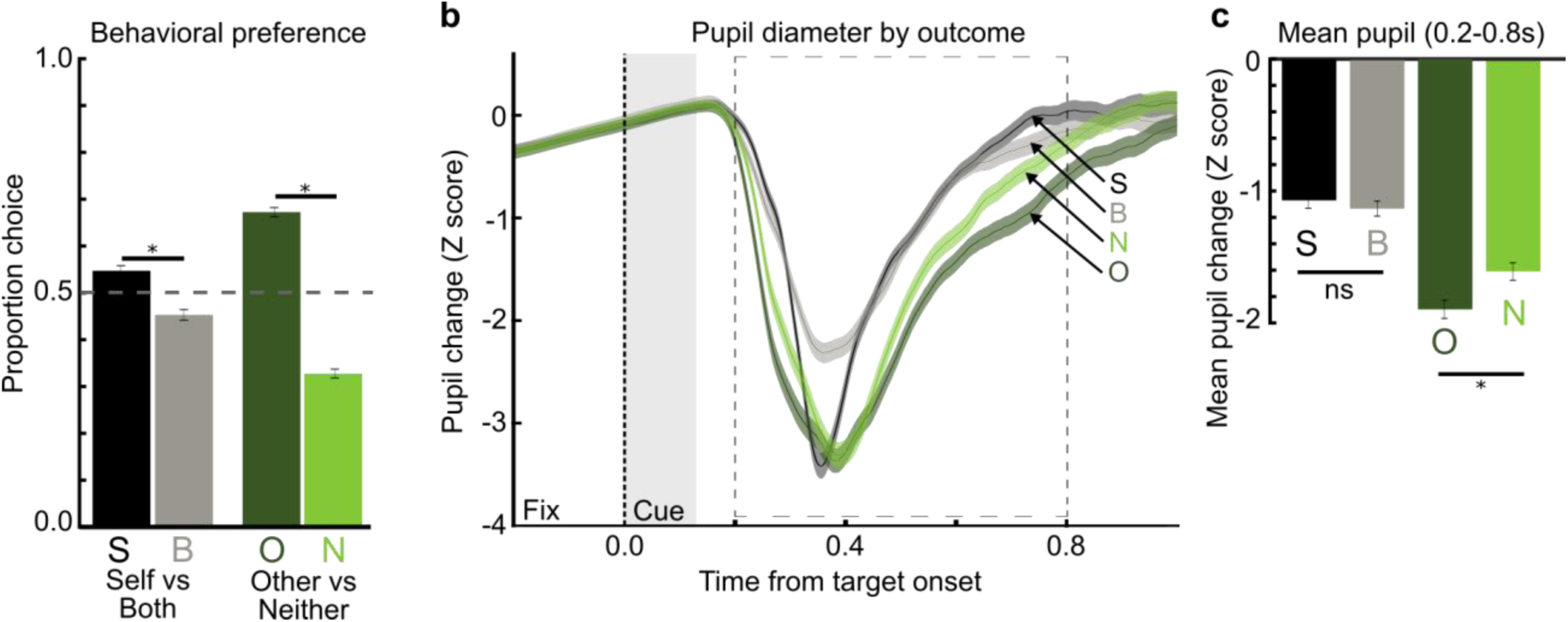
Pupils were more constricted in anticipation of preferred prosocial *other* trials than *neither* trials. a. Behavioral preference from choice trials. Actor monkeys chose between *self* and *both* reward conditions on one trial type and between *other* and *neither* reward conditions on another trial type. Proportion choice indicates decision preferences for choosing *self* and choosing *other* in each condition. b. Relative changes in pupil size after *self* cue (S), *both* cue (B), *other* cue (O), and *neither* cue (N) trials are shown aligned to the onset of the cue (Cue) with previous fixation noted (Fix). Dashed line box indicates analysis epoch. Shorter analysis epochs showed similar effects. Error bars and shaded bands are ±SEM. c. Average pupil diameter for each outcome during the 600-ms analysis epoch. The *neither* (N) reward outcome is associated with a larger pupil diameter than the *other* (O) reward outcome. Reward received trials (S, B) are associated with larger pupil diameter than reward forgone (O, N) trials.

Pupil size predominantly reflected the differences between the reward forgone (*other* and *neither* trials) and reward received (*self* and *both*) conditions (Fig 4b) as measured by one-way ANOVA with reward outcome as the factor (F_(3,40)_ = 4.15, p < 0.001). Within reward received trials, pupil diameter did not differ significantly between *self* (M ± SEM; −1.12 ± 0.06) and *both* (−1.07 ± 0.06) reward outcomes (Tukey test, p=0.93). This is not altogether unexpected given the strong role of primary reward in autonomic arousal and that monkeys were consuming a juice reward in both circumstances.

Importantly, as in Experiment 1, monkeys had larger pupil diameters following the *neither* cue (−1.7 ± 0.75) than the *other* cue (−1.9 ± 0.76, Tukey test p<0.01). This difference is notable as it is opposite the explicit social preference of the animals in which they preferred choosing *other* over *neither.* Taken together, the pattern of findings from Experiments 1 and 2 indicates that pupillary responses are not indexing prosocial preference.

## 4. General discussion

Across two laboratories, with different monkeys, different versions of a social reward allocation task, and different stimuli, we found that monkeys’ pupils were paradoxically narrower in anticipation of the preferred prosocial outcome (*other* trials) relative to the less preferred antisocial outcome (*neither* trials). This is contrary to what is usually observed in studies that manipulate reward magnitude, in which pupil size continually increases as outcomes become more preferred^10^. In this task, vicarious reward does not correspond with increasing pupil diameter.

One parsimonious explanation for this orthogonal ordering of outcome preference and pupil size is that trial preference indexes outcome valence, pupil size indexes outcome salience, and the relation between valence and salience is U-shaped (Fig 5). Under this explanation, *self* and *both* have a strong positive valence and high salience, *other* has a weak positive or even neutral valence and a low salience, and *neither* has a negative valence and a moderate salience. Evaluating this explanation will require additional studies, perhaps using different manipulations of outcome valence^2^.

**Figure 5.**
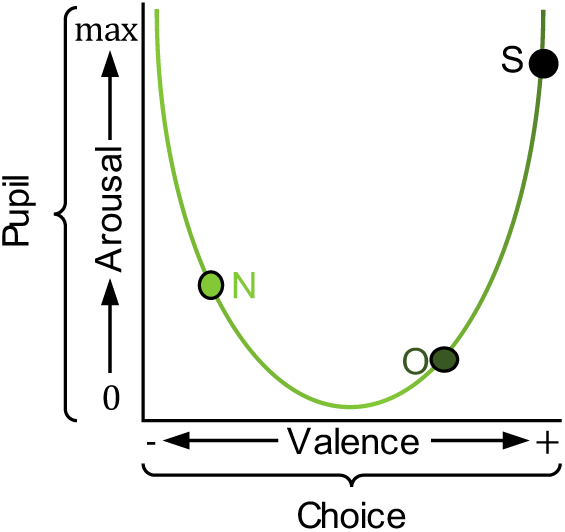
Hypothetical relation between arousal and valence. Outcome valence, from negative to positive, is depicted as a U-shaped function of the autonomic arousal produced by the outcome. Reward outcomes are placed in hypothetical locations along this continuum. Our results are consistent with the explanation that pupil size tracks arousal whereas outcome preference tracks valence. Reward to *Self* (S) would have high arousal and positive valence. Reward to *Other* (O) would have low arousal and positive valence. Reward to *Neither* (N) would have medium arousal and negative valence.

One alternative explanation is that the wider pupils in anticipation of *neither* rewards, relative to *other* rewards, reflects more effortful cognitive processing. In humans, pupils widen during problem solving and decision making, and this dilation is more pronounced when subjects are uncertain about their decision^17-19^. For our monkeys, it is possible that accepting a trial that would deny juice to their partner was cognitively effortful, involved more covert attention, or was done with uncertainty. However, comparing the pupil traces in the Social and Nonsocial session in Experiment 1 suggests that pupils were abnormally constricted on *other* trials rather than being abnormally dilated on *neither* trials. This “level-of-processing” hypothesis will require more investigation.

A second alternative explanation for the different orders of trial completion rates and pupil widths is that monkeys give juice to another monkey under duress. Wide pupils usually predict preferred outcomes, so the constricted pupils in anticipation of juice reward to the other monkey might indicate that actor monkeys found the prosocial choices to be aversive. Primates do engage in social interactions they find aversive, such as when subordinate macaques tolerate dominant monkeys stealing stored food directly out of their cheek pouches^20^. Such obligate prosociality is an intriguing hypothesis, but unlikely. Obligate prosociality should occur more in subordinate individuals but our effect was observed in both dominants and subordinates. Further, anecdotal evidence suggests that the prosocial preference for *other* over *neither* may even be stronger in dominant individuals who would have no need to oblige their subordinate partners ^1,2^. Lastly, although our actors knew their testing partners, they did not live together. Thus, it is unlikely that they grudgingly preferred the *other* rewards because they feared later retribution.

Our pupil size effect mirrors the group firing rate pattern of neurons on the gyral portion of the anterior cingulate cortex (ACCg) found in a previous study using cued social reward outcomes^7^. Individual ACCg cells were active in anticipation of reward delivery to *self, other*, or *both* monkeys. As a population, in the cued-reward condition, which is closest to the conditions used in this study, the ACCg neuronal firing rate was numerically highest to *self*, next highest to *neither*, and lowest to *other* (see Chang et al., 2013, Fig 3e). The same ordering of ACCg firing rate and pupil size serves as supporting evidence linking monkeys’ prosocial behavior, autonomic arousal, and ACCg activity.

The ACC is strongly connected to the locus coeruleus^21,22^, and locus coeruleus activity correlates with pupil size^14^. Causally, aspiration lesions of the subgenual ACC abolish the sustained pupil dilation in anticipation of reward^13^ and aspiration lesions of the ACC gyrus reduce the delay monkeys normally exhibit when retrieving food in the presence of social stimuli^23^. Together, these findings suggest that the relation between prosocial behavior and autonomic arousal relies on a network of brain regions including the ACC. Future research should examine the causal contributions of the ACC to monkeys’ prosocial tendencies and pupil size in this vicarious reinforcement test.

These findings, and the replicability and generalizability they demonstrate, suggest that the option of delivering juice rewards to no one instead of to the other individual in the social reward allocation task represents a particularly salient outcome for actor monkeys. Furthermore, these findings indicate that there is an interplay between reward and salience in the social reward allocation task, and likely in other social interaction paradigms. Lastly, these data demonstrate that autonomic measures like pupil size provide unique information that would not otherwise be detected via traditional measures like trial completion rates or choice preferences. Future studies of social cognition will benefit from including autonomic measures.

## Author Notes

We thank Drs. Andrew R. Mitz, Jaewon Hwang, and Vincent D. Costa for technical support and analysis advice. This work was supported by funding from the National Institute of Mental Health (NIMH grants R01 MH120081, R01 MH110750, and ZIA MH00288712).

